# Biomedical text corpora for multi-entity recognition in diet-related metabolic syndrome research

**DOI:** 10.1101/2025.09.04.673740

**Authors:** Antoine D. Lain, Stephanie Go, Alisha Mahmud, Shruti Rajendra, Ainara Cano, Katerina Loupasaki, Georgios Theodoridis, Maider Bizkarguenaga, Yajie Gu, Olga Deda, Ricardo Conde, Nieves Embade, Ángela de Diego, Nerea Burguera, Danai Rossiou, Rubén Gil-Redondo, Domniki Gallou, Itziar Tueros, Rakesh Velmurugan, Vasiliki Gkanali, Mercedes Caro, Petros Pousinis, George Alektoridis, Sara Arranz, Nasos Nikolopoulos, Xingchen Yan, Rebeca Fernández-Carrión, Thomas Rowlands, Donghee Choi, Marek Rei, Chris Cave-Ayland, Adrian D’Alessandro, Tim Beck, Joram M. Posma

## Abstract

We present here five biomedical, multi-entity corpora that can be used as benchmarks for named-entity recognition (NER), targeted to literature on metabolic syndrome. The CoDiet-Gold corpus contains annotations for 500 full-text publications and 348,406 annotations. It is divided into CoDiet-Gold-public (450 documents) and CoDiet-Gold-private (50 documents). Each document was independently annotated by two human experts, with disagreements fully adjudicated by a third expert. The CoDiet-Electrum corpus (2,998,273 annotations) contains 4,423 publications that were annotated using case-insensitive matching of the surface forms with punctuation ignored, found in CoDiet-Gold-public. Finally, for the same 4,423 documents, two fully machine-annotated corpora CoDiet-Bronze (2,938,738 annotations) and CoDiet-Silver (2,298,988 annotations), were created by utilising existing NER algorithms to annotate these. These corpora contain categories (organisms, disease, genes, proteins, metabolites) that add depth to existing corpora, as well as new categories that do not appear in other corpora (food, dietary methods, sample types, computational methods, study methodology, population characteristics, data types, and microbiome).

**Availability:** These corpora and annotations are available at Zenodo in various forms (https://doi.org/10.5281/zenodo.22097875, https://doi.org/10.5281/zenodo.22097879, and https://doi.org/10.5281/zenodo.22097881).

## Background & Summary

The expanding volume of scientific literature makes it increasingly difficult for researchers to remain up to date with emerging knowledge and to perform comprehensive literature review (1, 2). In biomedicine, computational approaches including information retrieval systems, natural language processing (NLP), and large language models (LLMs) are increasingly used to support literature exploration and evidence synthesis. However, extracting structured information from biomedical publications remains challenging because scientific articles contain specialised terminology, heterogeneous reporting styles, and domain-specific concepts that are often expressed across full-text documents rather than abstracts alone (3). These challenges motivate the creation of structured, machine-readable literature resources that support automated extraction, comparison, and integration of biomedical information.

The CoDiet corpora (4) were developed to support large-scale analysis of the scientific evidence linking diet and health. Understanding the relationship between food and disease remains challenging because dietary effects are influenced by many interacting factors, including genetics, lifestyle, metabolism, microbiome composition, and environmental exposures and new methods for dietary assessment are needed to deal with this on a population scale (5). At the same time, non-communicable diseases (NCDs), including metabolic disorders and cardiovascular diseases (CVDs), are the leading cause of death worldwide and their prevalence has increased substantially with, at present, CVDs contributing 29% of worldwide deaths and projected to reach 86% by 2050 (6). The CoDiet consortium was established to develop data-driven and artificial intelligence approaches to better understand the relationships between diet and health and improve the use of nutrition and health data. However, current evidence relevant to understanding these interactions is distributed across a rapidly growing and heterogeneous body of literature. The CoDiet corpus was created to provide a structured full-text resource that enables automated extraction and integration of dietary and biomedical information from scientific publications.

Training of models for biomedical named-entity recognition (NER) relies on the availability of biomedical text corpora in machine-readable format (7), however most corpora focus on a small number of annotation types (see Table 1). The annotation categories available from these corpora dictate the developments made in the field of bioNLP, and with newer corpora still focusing on the same categories as prior ones they add depth rather than breadth.

**Table 1.** Available biomedical corpora used for named-entity recognition. Short form, citation, year of release, text type (and number), number of total annotations, and number of categories of annotations are indicated.

| Name | Citation | Year | Sentences (n) | Annotations (n) | Categories (n) |
| --- | --- | --- | --- | --- | --- |
| BC1GM | (8) | 2004 | 10,000 | 12,000 | 1 |
| MedTag | (9) | 2005 | 25,965 | 45,000 | 2 |
| BC2GM | (10) | 2006 | 20,000 | 44,500 | 1 |
| BioInfer | (11) | 2007 | 1,100 | 6,349 | 3 |
| EBI-disease | (12) | 2008 | 856 | 908 | 1 |
| BioNLP09 | (13) | 2008 | 9,372 | 14,963 | 2 |
| BC8GPE | (14) | 2023 | 2,170 | 3,501 | 1 |
| Name | Citation | Year | abstracts (n) | Annotations (n) | Categories (n) |
| JNLPBA | (15) | 2003 | 2,000 | 59,963 | 5 |
| GENIA | (16) | 2003 | 2,000 | 93,293 | 8 |
| GETM | (17) | 2010 | 150 | 656 | 2 |
| DECA | (18) | 2010 | 644 | 6,407 | 2 |
| MetEnz | (19) | 2011 | 296 | 1,853 | 2 |
| PTM | (20) | 2011 | 360 | 4,698 | 1 |
| Ex-PTM | (20) | 2011 | 360 | 4,698 | 2 |
| MLEE | (21) | 2012 | 262 | 8,227 | 7 |
| BioNLP11EPI | (22) | 2012 | 1,200 | 15,811 | 2 |
| Species-800 | (23) | 2013 | 800 | 3,708 | 1 |
| Variome | (24) | 2013 | 10 | 5,650 | 4 |
| AnatEM | (25) | 2013 | 962 (250 sentences) | 13,701 | 4 |
| BioNLP13PC | (26) | 2013 | 525 | 15,901 | 5 |
| DDI | (27) | 2013 | 233 (792 sentences) | 18,502 | 1 |
| tmVar | (28) | 2013 | 500 | 1,431 | 1 |
| ChemDNER | (29) | 2013 | 10,000 | 84,355 | 2 |
| NCBI-disease | (30) | 2014 | 793 | 6,892 | 1 |
| BC5CDR | (31) | 2014 | 1,500 | 131,343 | 2 |
| LocText | (32) | 2015 | 54 | 201 | 1 |
| ChemDNER patents | (33) | 2015 | 21,000 | 99,634 | 2 |
| ME corpus | (34) | 2016 | 271 (2,288 sentences) | 4,411 | 2 |
| ChemProt | (35) | 2017 | 1,682 | 42,608 | 2 |
| OpenMinTeD-abs | (36) | 2018 | 200 | 5,582 | 5 |
| PGR | (37) | 2019 | 1,712 | 19,511 | 2 |
| DrugProt | (38) | 2020 | 4,250 | 108,387 | 2 |
| NLM-Gene | (39) | 2021 | 550 | 15,581 | 2 |
| DIMB-RE-abs | (40) | 2025 | 165 | 7,776 | 15 |
| Name | Citation | Year | full-texts (n) | Annotations (n) | Categories (n) |
| BC3IMT | (41) | 2009 | 2,590 | 5,664 | 3 |
| Linnaeus | (42) | 2010 | 100 | 4,077 | 1 |
| BioNLP11ID | (22) | 2012 | 30 | 11,082 | 4 |
| CRAFT | (43) | 2012 | 97 | 99,907 | 10 |
| BC4GO | (44) | 2013 | 200 | 10,898 | 2 |
| CellFinder | (45) | 2014 | 20 | 16,026 | 8 |
| ChemPatCorpus | (46) | 2014 | 47 | 35,337 | 4 |
| BRONCO | (47) | 2016 | 108 | 403 | 5 |
| OpenMinTeD-full | (36) | 2018 | 100 | 41,313 | 5 |
| GNI | (48) | 2018 | 499 | 88,629 | 5 |
| NLM-Chem | (49) | 2021 | 150 | 40,467 | 1 |
| EuropePMC | (50) | 2023 | 300 | 72,378 | 3 |
| DIMB-RE-res | (40) | 2025 | 30 | 6,674 | 15 |

In this article, we describe the creation of the CoDiet-Gold corpus, a machine-readable multi-entity corpus curated from biomedical full-text publications using a combined dictionary-, regular expression-, and deep learning (DL)-based annotation workflow followed by experts manually, and independently, annotating documents. The resource consists of a manually reviewed gold-standard corpus of 500 full-text publications, in which each document was independently annotated by two human annotators and disagreements, including annotations produced by only one annotator, underwent independent adjudication. The CoDiet-Gold corpus is split into a public version of 450 documents with all annotations shared (4), and a private version of 50 documents whose annotations are kept separate to prevent data leakage. In addition, we release three corpora generated from 4,423 full-text Open Access publications with redistribution-permissive licensing (Creative Commons or similar license) (51). The machine-annotated corpora are CoDiet-Bronze (all machine-generated annotations retained), CoDiet-Silver (machine annotations after overlap resolution and post-processing), and CoDiet-Electrum (annotations generated using the validated labels and vocabulary derived from CoDiet-Gold-public). Annotations are available for all 4,423 articles (4), and a subset of 3,965 documents with licenses that permit sharing of derived work is provided fully processed (52). Together, these resources provide structured full-text annotations and accompanying metadata in a machine-readable format for reuse and evaluation. Additionally, we provide code that allows users to process all documents from their original formats into the final BioC-JSON annotated documents to allows the replication of the entire corpus, including all documents with licenses that prohibited sharing derived work (i.e. converting from XML to BioC-JSON, adding in annotations to BioC-JSON files).

## Methods

### Search Query

A search query to find relevant publications was built up from a two-day collaborative exercise during the kick-off meeting of the CoDiet project to reach a consortium-wide consensus on the scope of the literature review, specifically on data entities and phenotypes of interest, in January 2023. This was run using a Mentimeter survey to gather responses on 1) disease, conditions, and phenotypes, 2) data, methodologies, and methods, 3) sample types, 4) foods, diets, and diet methodologies, and 5) exclusion terms. 26 CoDiet collaborators contributed 1,043 individual terms deemed relevant, with 48 exclusion terms mentioned. A consensus discussion around the summary of 1,043 terms resulted in an agreement to focus on metabolic syndrome and its components with multi-modal data relating to various omics and wearable technologies. This was subsequently refined into six categories (A-F) of terms to be searched in titles and abstracts: A) 121 disease/phenotype terms (e.g. cardiometabolic syndrome, dyslipidemia, glucose response), B) 153 diet-related terms (e.g. caloric restriction, diet diaries, nutritional behaviour), C) 23 data types (e.g. image, urine, stool), D) 90 methodologies (amplicon sequencing, camera technology, polygenic risk score), E) 81 study types (e.g. cohort study, randomized controlled clinical trial, personalised nutrition), and F) 49 exclusion terms (e.g. cancer, NAFLD, saliva). A table with all inclusion and exclusion terms is provided as Supplementary Material. These terms were used to create search queries for Web of Science (WoS), PubMed and PubMed Central (PMC) of the form of [any term from A] AND [any term from B] AND [any term from C] AND [any term from D] AND [any term from E] NOT [any term from F]. WoS, PubMed, and PMC were searched manually on 07 July 2023 and lists of digital identifiers (WoS ID, PMID, PMCID, DOI) were extracted. A total of 3,372 identifiers from PubMed, 1,645 from PMC and 12,687 from WoS were found as potentially relevant to CoDiet.

### Data – Publications

We used application programming interfaces (APIs) to resolve the PMC and WoS identifiers to PubMed identifiers (PMIDs), resulting in 12,112 PMIDs. These identifiers were used as input to CADMUS (53), an automated full-text retrieval system for large-scale biomedical corpus generation. CADMUS integrates multiple retrieval strategies, including publisher websites, PubMed Central (PMC), DOI resolution services, and institutional subscription access, enabling retrieval of both Open Access (OA) publications and articles available through host institution licences. Retrieval quality is controlled through internal validation procedures based on publication metadata and document characteristics, and previous evaluation of CADMUS demonstrated high fidelity of retrieved full texts when compared with corresponding versions available through PMC. Publications that prohibit redistribution were separated from OA publications with redistribution-permissive licences (Creative Commons or similar license). The retrieved full-text publications were converted from their native formats (HTML and XML) into the machine-readable BioC-JSON format (54) using Auto-CORPus (55), with document sections standardised and mapped to Information Artifact Ontology (IAO) (56) terms. The released corpora are distributed in various formats to conform with all individual licenses.

All redistributable documents in the CoDiet corpus are shared in their original formats (51) and separately the annotations as single JSON files in BioC format for each document (4). Fully processed documents (that permit derivations) are shared as BioC-JSON files and are represented as structured dictionaries containing publication metadata, passage-level text segmentation, annotations, normalised identifiers, and annotation metadata. To support transparency and reproducibility, each BioC document additionally stores provenance information in the `infons` dictionary at the document level, including the original source from which the publication was retrieved (e.g. publisher, PMC, DOI resolution), the retrieval link where available, and associated meta-data generated during document acquisition and processing (including license terms). For CoDiet, CADMUS retrieved 10,173 publications out of 12,112 (83.99%) directly. An additional 251 publications were identified through PMC and 740 through the doi.org API, resulting in a final corpus of 11,164 publications (92.17% retrieval rate). As part of the wider CoDiet research project, non-OA publications were retrieved that were available through Imperial College London subscriptions, however these are not shared or redistributed due to copyright laws and therefore are not part of the data described in this article. The re-distributable subset comprised 3,688 full-text publications and forms the basis of the dataset presented in this work. Following document standardisation, passages without relevant scientific content for annotation, including ‘Author Information’ and ‘’ were removed in the final BioC-JSON versions.

### Annotation Software and Process

From the 3,688 publications, 500 were chosen at random (spanning the entire time period data was available for to avoid recency bias) to be annotated by human experts. We recruited annotators from within the CoDiet consortium via an open call for recruitment during a consortium-wide meeting in October 2023 and via email. Additionally, we recruited further volunteers from within one of the institutions (Imperial). A total of 44 people initially volunteered from five partners (Imperial, AUTh, AZTI, CICBIO, UVEG). We ran a dedicated online meeting for all potential annotators to demonstrate the collaborative software (see below) used for the annotation process. During this meeting the participants were briefed on the category types and were given examples for each of these 13 categories (see below). The meeting was recorded for the other volunteers and shared with them after the session.

TeamTat (57) is a collaborative, online text annotation tool, designed to facilitate the annotation process for large document collections to ready these for downstream tasks such as training NLP models. It has a user-friendly interface for both project managers and annotators enabling efficient labelling. Anonymity features are available to better reduce bias, while quality control is performed after annotations are done to ensure data accuracy. In our implementation of TeamTat, we added a customisable visibility tab to help annotators focus on specific categories they have expertise in. Our customised version of the TeamTat software was deployed by hosting it on an Azure cloud platform to allow (credentialised) access for any CoDiet member.

After the demo meeting we reached out to all volunteers to ask them for their self-perceived expertise for the 13 categories, their time commitment (number of articles per week/in total) and assigned articles in batches to the 38 volunteers that responded. Batches were released weekly and assigned semi-randomly, i.e. annotators were assigned articles that had entities relevant to their expertise as not all articles had all categories but were then assigned their specified number of articles for each batch at random. Three annotators that initially verified their involvement dropped out after having been assigned articles. Once an annotation round is started, TeamTat does not allow re-assigning articles to annotators (based on their user-id), hence we updated the login details of each user-id to give other annotators a secondary login to complete these articles. Over the course of the first 6 weeks of the task, we scheduled 1-2 weekly drop-in sessions via Microsoft Teams that any annotator can join and share their screen to get help/advice. These sessions were initially attended by 15-20 annotators each time, but over time (as instructions were clarified and annotators gained experience) this was reduced to 1-2 individuals. As a result of these drop-in sessions we updated the materials on the instructions and added further examples to a live Google Doc FAQ to help the annotators. A total of 12 batches were released until the end of January 2024. An archive of the final version of the FAQ is shared on Zenodo (4).

### Categories for Annotation

To provide a comprehensive coverage of biomedical concepts relevant to our target domain, we designed an annotation scheme encompassing 13 distinct entity categories (Table 2). These categories were chosen to capture biological and experimental information at multiple levels, ranging from molecular entities such as genes, proteins, and metabolites to broader study-level descriptors like methodology, sample types, and population characteristics. By including both biomedical concepts (e.g. microbial taxa, disease phenotypes) and contextual information (e.g. computational approaches, dietary assessment methods), this schema was used to capture a multi-faceted representation of the literature.

**Table 2.** Summary of the annotated categories. Descriptions of categories of annotations in the corpus with descriptions and examples given to annotators.

| Category | Description | Examples |
| --- | --- | --- |
| dietMethod | A broad description of the type of methodology used to capture dietary/nutrition data | food frequency questionnaire, nutritional epidemiology, dietary recommendations, personalised nutrition, dietary assessment |
| foodRelated | Food, nutrients, and diets that are mentioned in the context of intake, biomarkers measured in biofluids should not be included | Mediterranean diet, dietary fibre, high-fat diet, omega-3, apple |
| metabolite | Small molecules measured in biofluids and tissue, includes both metabolites and lipids | sphingomyelin (d18:0/17:0), Trimethylamine N-oxide, cholesterol, PC(34:0), glucose |
| microbiome | Microbial entities from (super)kingdoms archaea, bacteria and fungi, this can be of any taxonomic rank | *Methanobrevibacter smithii*, *Akkermansia* spp., *Candida albicans*, *Firmicutes*, *E. coli* |
| proteinEnzyme | Protein and enzyme mentions including protein hormones and lipoproteins, this is distinct from the gene mentions although these can overlap depending on context | alanine aminotransferase, LDL-cholesterol, oxidoreductase, glucagon, CRP |
| geneSNP | Gene and single-nucleotide polymorphism mentions, this can include both abbreviations for genes as well as full names | lipoprotein lipase gene, rs1571960363, *CRP* gene, *MALAT1*, *GNPAT* |
| diseasePhenotype | Any term related to a type of disease/phenotype/medical condition that is being studied or for which the results are relevant, we are primarily interested in entities around metabolic syndrome, however any non-communicable disease entity is also of interest | insulin resistance, atherosclerosis, prehypertension, obesity, stroke |
| sampleType | The sample types from which data is collected | 24hr urine collections, fasted, stool, video, gut |
| dataType | The type of data or technology used by the study to analyse/measure the samples | 16s rRNA gene sequencing, micro-camera assessment, bioimpedance, lipidomic, HPLC-MS |
| methodology | A broad description of the type of general study methodology used, this is different from the diet methodology | randomised controlled clinical trial, public health policy, personal monitoring, free-living cohort, cross sectional |
| modelOrganism | Descriptions of the type of organism the study is conducted with, i.e. human or animal data, this should not include microbiota | *Rattus norvegicus*, *C. elegans*, patient, human, mouse |
| populationCharacteristic | A broad range of descriptors from the cohorts or populations that are studied | physical activity, family history, lifestyle, female, age |
| computational | Computational/statistical methods used to analyse the data | logistic regression, machine learning, paired t-test, correlation, AI model |

We introduced two additional categories to assist annotators when they were uncertain about the appropriate category (unsureCoDiet) or identified potentially relevant entity types that could not be attributed to any of the predefined categories (potential). In total, 578 annotations were assigned the unsureCoDiet category, and 104 were assigned the potential category. All 682 annotations were reviewed independently by two adjudicators, regardless of whether they had already been annotated by other annotators. Following adjudication, 166 annotations were removed because they did not fit any of the defined categories, while the remaining annotations were assigned to their corresponding categories following agreement by two adjudicators. Therefore, no annotations labelled unsureCoDiet or potential remained in the final corpus.

### Annotation Pipeline

In order to help the annotators, we pre-filled the 500 documents with machine-generated annotations. We used a hierarchical system to decide on which annotation to use in case of disagreements between methods. In this system, we first aggregated annotations with identical spans and biomedical types. This involves consolidating identifiers and methods to maintain a single, aggregated annotation with comprehensive metadata. In cases where multiple annotations share the same span but differ in biomedical types, a rule-based approach is implemented. The rule-based method follows a priority order, favouring dictionary methods (see Table 3) populated with terms confidently identified with their biomedical types. Subsequently, DL methods were considered in a specific sequence (PhenoBERT (58), microbELP (59), TABoLiSTM (60), eNzymER (61), BERN2 (62)). After this, dictionary matching takes precedence over ontology matching (curated OBO ontologies), with the Metamap (63) ontology considered last due to the large numbers of potential false positives being annotated. When an annotation is encompassed by another (i.e. one is a complete subset of another), we retained the longest annotation. For overlapping annotations with distinct biomedical types, the rule-based decision approach is again invoked. However, if overlapping annotations share the same entity type (category), then they are merged to form a more extensive annotation that encapsulates all relevant information.

**Table 3.** Summary of the methods used to annotate entities for each of the 13 categories. Abbreviations: DB - dictionary-based annotation and normalisation, DL - deep learning model, RegEx - regular expression.

| Category name | Dictionaries, ontologies and algorithms used for annotation |
| --- | --- |
| dietMethod | DB (CoDiet: 105 terms, OBO ontology (one)) |
| foodRelated | DB (CoDiet: 44 terms, OBO ontology (cdno, foodon), UMLS ontology (food)) |
| metabolite | DB (HMDB (64) and LIPID MAPS (65) ontologies, 1,934,277 terms), DL (TABoLiSTM (60)) |
| microbiome | DB (NCBI taxonomy: 124,706 terms), DL (microbELP (59)) |
| proteinEnzyme | DL (BERN2 (62), eNzymER (61)) |
| geneSNP | DL (BERN2 (62)), RegEx (for SNPs) |
| diseasePhenotype | DB (CoDiet: 228 terms), DL (BERN2 (62), PhenoBERT (58)) |
| sampleType | DB (CoDiet: 24 terms, UMLS ontology (bodysubstance)) |
| dataType | DB (CoDiet: 92 terms, UMLS ontology (laboratoryprocedure)) |
| methodology | DB (CoDiet: 136 terms, UMLS ontology (researchactivity)) |
| modelOrganism | DB (CoDiet: 99 terms), DL (BERN2 (62)) |
| populationCharacteristic | DB (CoDiet: 39 terms) |
| computational | DB (CoDiet: 44 terms, OBO ontology (stato,obcs)) |

This structured approach was used to resolve discrepancies, combining the strengths of various methods while maintaining consistency and accuracy.

After performing NER across the 13 pre-defined categories, we normalised all identified entities by assigning a common identifier to similar entities, such as synonyms (e.g., “high blood pressure” and “hypertension”). Our normalisation process is based on the methods used during the NER stage. Entities identified with dictionary, ontology, and regular expression methods are normalised based on textual similarity. We used fuzzy matching to assign the identifier of the closest term from our database, accounting for differences using Levenshtein distance, which considers the number of inserted, deleted, or substituted characters. For entities identified via DL models, we employed contextual similarity obtained from their embedding values. These deep learning models, trained on large corpora, learn relationships between words by examining their co-occurrence within sentences. Through word embeddings, extracted from our newly trained models, where words are projected into a higher-dimensional space, the model can capture semantic similarities. By computing the distance between word vectors using the cosine similarity, we identify the closest known term from our curated database (e.g. for metabolites we created a combined Human Metabolome DataBase (HMDB) (64) and LIPID MAPS (65) dictionary of identifiers with names/synonyms) and assign its identifier to the newly identified entity.

The application of all methods to annotate all 3,688 OA documents, without resolving overlaps, resulted in a ‘bronze corpus’ with all machine-made annotations. Overlaps were resolved according to the rule-based scheme to prioritise the output of certain models over others, the result from this process resulted in the ‘silver corpus’ (see Figure 1). The annotations for these corpora (CoDiet-Bronze, CoDiet-Silver) have identifiers from the ontologies/methods they were annotated by.

**Fig. 1.**
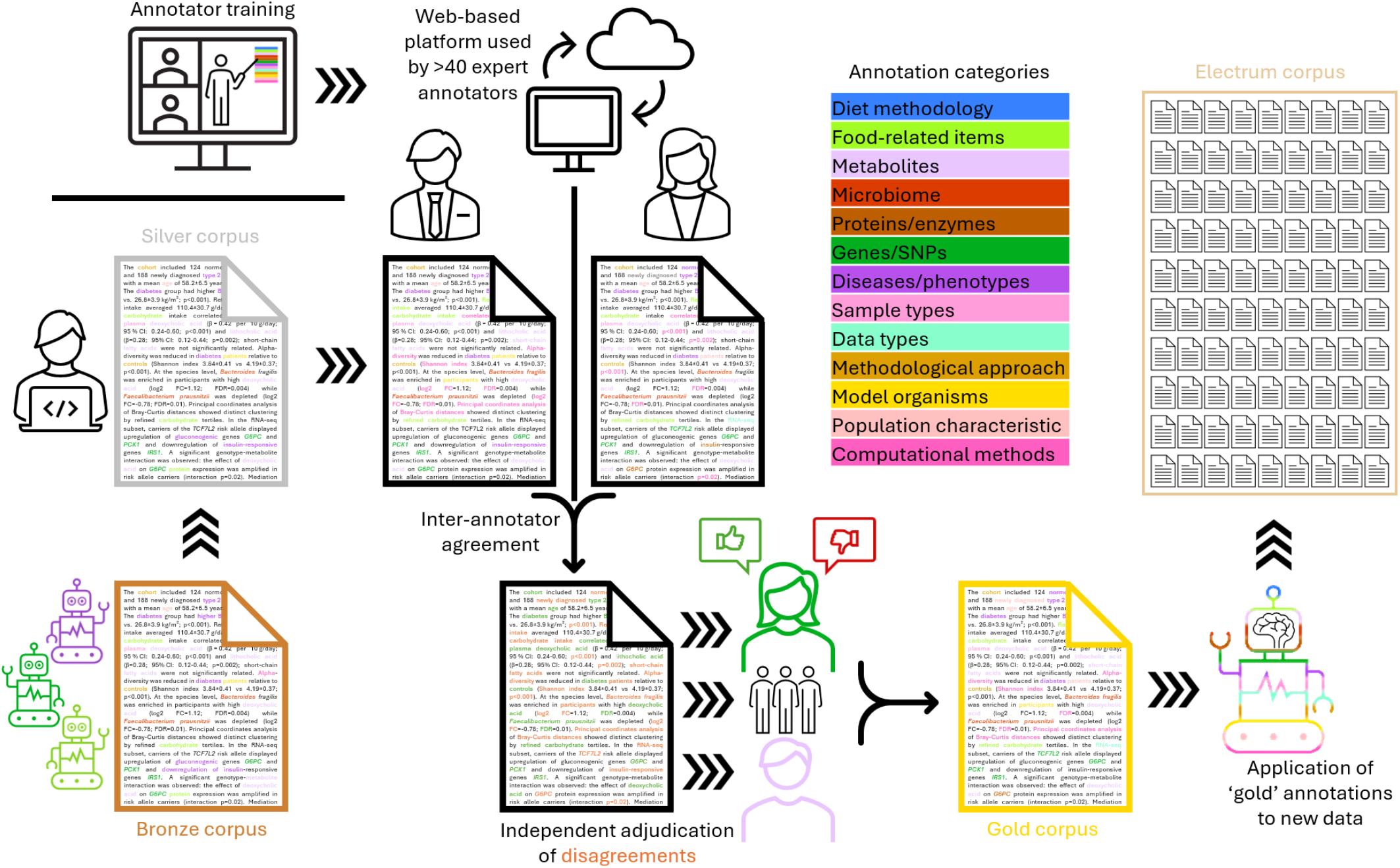
Overview of the process to curate a gold-standard text corpus. 500 Open Access full-text documents were processed using Auto-CORPus (55) and annotated using category/domain specific NER tools (bronze) with the majority of overlaps were resolved by means of a ranking of algorithms to take priority over another (silver). Over 40 annotators were trained to use the TeamTat annotation system (57) hosted on an Azure cloud server, and the 500 documents were assigned to annotators based on availability each week and expertise (e.g. articles without gene mentions were not commonly assigned to those with genetics expertise). Two annotators independently checked the machine annotations and added their own annotation in each document. The annotations were merged and disagreements adjudicated by an independent team of experts with domain expertise on the full set of 500 documents. The gold-standard corpus was then used to create a dictionary of entities to be annotated in a larger set of documents (‘electrum’ standard: human verified (gold) labels annotated using automated (silver) means).

### Data Record

The CoDiet data resources are publicly available through three Zenodo releases that separate the original source documents, annotation information, and directly distributed processed corpora. This structure supports reproducibility while preserving the original Open Access PubMed Central XML files separately from the generated annotated BioC.

The source code required to process and reconstruct the datasets is available through the *CoDietZenodo* GitHub repository (https://github.com/omicsNLP/CoDietZenodo) and is archived on Zenodo (https://doi.org/10.5281/zenodo.22142400). The workflow converts the original XML documents into BioC JSON format and incorporates the corresponding CoDiet annotations into the generated BioC documents. The original corpus generation and annotation pipelines are available through the *CoDietCorpus* GitHub repository (https://github.com/omicsNLP/CoDietCorpus) and are archived on Zenodo (https://doi.org/10.5281/zenodo.20643594). Benchmarking infrastructure and evaluation instructions for the CoDiet-Gold-private corpus are available through Codabench (https://www.codabench.org/competitions/11676/).

### CoDiet corpus – raw XML

The *CoDiet corpus (v*.*0*.*1) – raw XML* release is available through Zenodo (51). It contains unchanged XML files obtained from Open Access PubMed Central and constitutes the source material used for generating the CoDiet corpora. The files are distributed in their original XML form and are not modified within this release.

The release contains three compressed archives:

### CoDiet-Bronze-Silver-Electrum-XML.zip

4,423 XML documents corresponding to the publications used to generate the CoDiet-Bronze, CoDiet-Silver, and CoDiet-Electrum corpora.

### CoDiet-Gold-public-XML.zip

450 XML documents corresponding to the CoDiet-Gold-public corpus.

### CoDiet-Gold-private-XML.zip

50 XML documents corresponding to the CoDiet-Gold-private corpus.

The raw XML corpus is distributed under the most restrictive licence represented in the corpus, CC BY-NC-ND. The source documents remain unchanged, and the original licensing conditions associated with the individual source publications should be considered when using the corpus.

### CoDiet corpus – annotations

The *CoDiet corpus (v*.*1*.*0) – annotations* release is available through Zenodo (4) under a CC BY-NC 4.0 licence. This release contains annotation information separately from the source XML documents and consists of four compressed archives:

### CoDiet-Bronze-annotations.zip

annotation records for 4,423 documents.

### CoDiet-Silver-annotations.zip

annotation records for 4,423 documents.

### CoDiet-Electrum-annotations.zip

annotation records for 4,423 documents.

### CoDiet-Gold-public-annotations.zip

annotation records for 450 documents.

The annotation files are provided in BioC JSON format and contain the information required to reconstruct the CoDiet annotations. The annotation information can be incorporated into BioC documents generated from the corresponding raw XML files using the *CoDietZenodo* workflow.

This separation allows users to obtain the original XML source documents and the corresponding annotation information independently and to recreate the annotated BioC representations locally.

### CoDiet corpus – full-texts annotated

The *CoDiet corpus (v*.*1*.*1) – full-texts annotated* release is available through Zenodo (52). This release contains directly distributed processed BioC JSON files for documents that can be provided as pre-combined full-text and annotation representations under the applicable licensing conditions. The release is distributed under CC BY-NC-SA.

It contains five compressed archives:

### CoDiet-Bronze.zip

3,965 machine-annotated BioC JSON documents.

### CoDiet-Silver.zip

3,965 machine-annotated BioC JSON documents.

### CoDiet-Electrum.zip

3,965 machine-annotated BioC JSON documents.

### CoDiet-Gold-public.zip

409 BioC JSON documents containing human annotations.

### CoDiet-Gold-private.zip

46 unannotated BioC JSON documents.

Documents not included in the pre-combined full-text release can nevertheless be processed and reconstructed locally using the raw XML release, the annotation release, and the *CoDietZenodo* workflow (https://doi.org/10.5281/zenodo.22142400). This enables users to recreate the corresponding CoDiet datasets.

### CoDiet corpus

The CoDiet corpus has five corpora. **CoDiet-Gold-public** consists of 450 full-text documents independently annotated by two human annotators and adjudicated through additional independent review rounds until retained annotations had agreement from at least two human annotators. A subset of 409 pre-combined annotated BioC documents is provided in the full-text release.

**CoDiet-Gold-private** consists of 50 full-text documents reserved for benchmarking and evaluation through Codabench. The corpus is distributed without annotation labels. Evaluation labels associated with this corpus are not publicly released. The raw XML release contains 50 source documents, while the processed full-text release contains 46 unannotated BioC documents.

**CoDiet-Electrum** consists of 4,423 documents automatically annotated using the validated labels and vocabulary derived from CoDiet-Gold-public, with post-processing rules applied. The pre-combined full-text release contains 3,965 annotated BioC documents.

**CoDiet-Silver** consists of 4,423 documents automatically annotated using existing deep learning, rule-based, and dictionary-based approaches with overlap resolution and post-processing. The pre-combined full-text release contains 3,965 annotated BioC documents.

**CoDiet-Bronze** consists of 4,423 documents automatically annotated using existing deep learning, rule-based, and dictionary-based approaches without post-processing. The pre-combined full-text release contains 3,965 annotated BioC documents.

### Data format and reconstruction

The raw source documents are provided as XML files. The *CoDietZenodo* workflow (https://doi.org/10.5281/zenodo.22142400) converts these XML files into UTF-8 encoded BioC JSON documents. The resulting documents contain structured text, metadata, and the BioC document structure required for incorporating CoDiet annotations.

The annotation release contains the information required to populate the corresponding BioC documents with annotations. Annotation records include text span information, character offsets, location information, and entity category labels. Finally, for the Bronze and Silver corpora, normalised identifiers were provided directly by the *CoDietCorpus* pipeline and linked to external ontologies. In contrast, identifiers for the Electrum and Gold-public corpora were obtained through case-insensitive matching of surface forms, after replacing spaces (‘ ‘) and commas (‘,’) with underscores (‘_’).

Using the *CoDietZenodo* workflow, users can first convert the corresponding raw XML documents into BioC JSON format and subsequently incorporate the matching annotations from the annotation release. This workflow enables reconstruction of the complete CoDiet datasets from the separate source XML and annotation resources.

The *CoDiet corpus (v*.*1*.*1) – full-texts annotated* (52) provides ready-to-use BioC JSON files for the subset of documents distributed directly as processed full-text representations. Users requiring the complete corresponding corpus can instead use the raw XML and annotation releases together with the provided reconstruction code.

Each document is identified using its corresponding PubMed Central identifier (PMCID), named as {PMCID}.{file extension}, enabling files from the raw XML and annotation releases to be matched during corpus reconstruction.

### Data Overview

The CoDiet resource consists of five corpora: CoDiet-Bronze, CoDiet-Silver, CoDiet-Electrum, CoDiet-Gold-public, and CoDiet-Gold-private. The corpora differ in their annotation methodology and intended use, and are distributed through releases containing source XML files, annotation information, and pre-combined BioC documents.

Table 4 provides an overview of the five corpora and their availability across the three components of the CoDiet data resource. The source XML column indicates the number of unchanged XML documents available from Open Access PubMed Central. The annotation files column indicates the number of documents for which CoDiet annotation information is available separately. The pre-combined BioC column indicates the number of ready-to-use processed BioC JSON documents containing the corresponding full text and annotations.

**Table 4.** Overview of the CoDiet corpora and their availability across the three components of the data resource. Source XML files are distributed unchanged. Annotation files are distributed separately from the source documents. Pre-combined BioC files provide ready-to-use processed full-text documents containing annotations.

| Corpus | Annotation type | Source XML | Annotation files | Pre-combined BioC |
| --- | --- | --- | --- | --- |
| CoDiet-Gold-public | Human | 450 | 450 | 409 |
| CoDiet-Gold-private | None publicly released | 50 | – | 46 |
| CoDiet-Electrum | Machine | 4,423 | 4,423 | 3,965 |
| CoDiet-Silver | Machine | 4,423 | 4,423 | 3,965 |
| CoDiet-Bronze | Machine | 4,423 | 4,423 | 3,965 |

The number of pre-combined BioC documents differs from the number of source XML and annotation files because of copyright restrictions. The complete corresponding datasets can be reconstructed from the unchanged source XML files and the separately distributed annotations using the *CoDietZenodo* workflow. CoDiet-Gold-public contains independently annotated and adjudicated human annotations and is intended for model training. CoDiet-Gold-private is reserved for benchmarking through Codabench, and its evaluation annotations are not publicly released. CoDiet-Bronze, CoDiet-Silver, and CoDiet-Electrum contain machine-generated annotations produced using different automated annotation and processing strategies.

The distribution of annotations across the 13 entity categories is presented in Table 5. For each category and corpus, the table reports the total number of annotations and the number of unique annotated surface forms with punctuation ignored. Unique surface forms were calculated using case-insensitive matching with punctuation ignored. The CoDiet-Gold-private is not reported by entity category because its annotations are reserved for benchmarking and evaluation. The Bronze corpus contains annotations generated using automated deep learning, rule-based, and dictionary-based approaches without post-processing. The Silver corpus uses corresponding automated approaches with overlap resolution and post-processing. The Electrum corpus was generated using the validated labels and vocabulary to perform case-insensitive matching of the surface forms with punctuation ignored, found in CoDiet-Gold-public. CoDiet-Gold-public consists of human annotations independently annotated and adjudicated through additional review rounds.

**Table 5.** Summary of Bronze, Silver, and Electrum (n = 4,423), and Gold-public (n = 450) CoDiet corpora. The asterisk (*) indicates the addition of 392,087 annotations overlapping multiple categories that were not assigned to a single category. The CoDiet-Gold-private corpus is not reported per category to prevent reverse engineering for prediction optimisation (total 33,991, on average 680 per document). The unique was counted using case-insensitive matching of the surface forms with punctuation ignored.

| Category | Bronze |  | Silver |  | Electrum |  | Gold-public |  |
| --- | --- | --- | --- | --- | --- | --- | --- | --- |
|  | Total | Unique | Total | Unique | Total | Unique | Total | Unique |
| dietMethod | 103,629 | 85 | 83,962 | 83 | 16,384 | 367 | 1,851 | 351 |
| foodRelated | 577,156 | 6,226 | 466,734 | 5,657 | 435,970 | 9,590 | 55,783 | 5,720 |
| metabolite | 285,859 | 16,428 | 207,087 | 14,850 | 173,918 | 3,818 | 38,005 | 3,641 |
| microbiome | 99,222 | 3,944 | 81,351 | 3,516 | 156,784 | 1,200 | 13,868 | 962 |
| proteinEnzyme | 16,497 | 1,485 | 13,168 | 1,232 | 141,001 | 2,598 | 19,726 | 2,756 |
| geneSNP | 157,904 | 31,371 | 145,767 | 29,979 | 19,956 | 1,296 | 5,402 | 1,558 |
| diseasePhenotype | 573,457 | 67,388 | 365,127 | 59,433 | 559,848 | 6,378 | 64,506 | 4,792 |
| sampleType | 215,819 | 2,607 | 185,372 | 2,321 | 263,981 | 1,775 | 28,007 | 1,175 |
| dataType | 172,211 | 7,270 | 163,658 | 6,068 | 161,506 | 2,913 | 17,614 | 2,297 |
| methodology | 151,905 | 6,679 | 142,941 | 6,006 | 92,275 | 1,253 | 8,605 | 852 |
| modelOrganism | 459,228 | 11,197 | 318,481 | 10,478 | 251,084 | 665 | 28,440 | 601 |
| populationCharacteristic | 100,644 | 32 | 100,584 | 32 | 214,580 | 1,562 | 18,427 | 658 |
| computational | 25,207 | 135 | 24,756 | 135 | 118,899 | 2,171 | 14,181 | 1,787 |
| Total: | 2,938,738 | 154,847 | 2,298,988 | 139,790 | 2,998,273* | 35,586 | 314,415 | 27,150 |
| average per article | 664 |  | 520 |  | 678 |  | 699 |  |

### Technical Validation

The quality of the CoDiet-Gold corpus (4) was assessed through independent annotation and multi-stage adjudication. Each of the 500 selected full-text publications was independently annotated by two human experts. Annotators could access the pre-filled machine annotations but were blinded to annotations produced by the paired expert to minimise confirmation bias.

Inter-annotator agreement (IAA) was measured using the F1-score computed over annotation spans and assigned categories. Across all categories, the average IAA was 0.78. Agreement varied between categories, with the highest agreement observed for `proteinEnzyme` (F1 = 0.86) and the lowest for `dataType` (F1 = 0.67).

To construct the final gold-standard corpus, annotations from both annotators were merged and all disagreements, including annotations created by only one annotator, were reviewed through independent adjudication. Five adjudicators participated in this process and difficult cases underwent additional review rounds until each annotation retained in the final corpus had agreement from at least two human annotators. This process ensured that annotations included in the released gold corpus were supported by human consensus rather than a single annotator decision.

### Annotation Guidelines and Evaluation Criteria

No standalone annotation guideline document was created for this corpus. Instead, annotation guidance was developed iteratively throughout the annotation process and distributed to annotators through email communications, live discussion sessions, and updates to a shared FAQ document (4). Annotators received an initial training session introducing the annotation categories, example annotations, and use of the annotation platform. During the annotation campaign, batches of documents were released progressively and guidance was updated based on questions raised by annotators and observations from the adjudication process. In addition, adjudicators met regularly (typically 3-5 times per week during active annotation periods) to discuss difficult cases, refine annotation decisions, and harmonise annotation practices across categories. Outcomes from these discussions were communicated back to annotators through follow-up meetings, town-hall sessions, and email updates.

Annotation decisions followed a set of practical conventions. Nested and overlapping annotations were not permitted and annotators were instructed to select a single span representing the most appropriate entity mention. Annotation spans aimed to capture the full semantic meaning of an entity while avoiding unnecessary modifiers; descriptive adjectives were excluded unless required to preserve the entity meaning (e.g. “high blood pressure” was annotated as a complete entity). Annotators were instructed to annotate textual variations and synonymous expressions as they appeared in the original text where these referred to concepts included in the annotation schema. For example, where sample collection conditions referred to fasting status, all equivalent textual expressions describing fasting were annotated under the corresponding category.

IAA was calculated using exact agreement criteria. Two annotations were considered matching only when the annotated text span, character offsets, and assigned category were identical. Partial span overlap and disagreements in category assignment, including semantically related categories (e.g. gene versus protein), were counted as disagreements and treated as incorrect matches during IAA calculation.

### Creation of Gold Standard Corpus

The annotators reviewed the full-text articles independently, i.e. human annotators could only access the pre-filled machine annotations and not annotations produced by the other paired annotator. The human annotator-reviewed machine annotations were combined with new annotations created during manual annotation process.

Annotations from both annotators were merged and all disagreements, including annotations produced by only one annotator, were submitted for independent adjudication. Five adjudicators participated in the review process, and additional review rounds were performed where required until each annotation retained in the final corpus had agreement from at least two adjudicators.

This process was applied to the complete set of 500 full-text articles to generate a fully manually reviewed gold-standard corpus (Figure 1). The resulting corpus was split into 450 articles (314,415 annotations) for public release as training and validation data (90%) and 50 articles (33,991 annotations; 10% of documents) to remain hidden on Codabench for benchmarking and evaluation.

Quality assessment of the annotation process, including inter-annotator agreement and adjudication outcomes, is described in the Technical Validation section above.

### Electrum Standard Corpus

To expand coverage beyond the original collection, an additional 1,342 documents published between 1 January 2023 and 30 September 2025 were identified through an updated literature search performed on 14 October 2025. Of these, 757 publications met the redistribution criteria and were not already included in the initial set of 3,688 documents. Following retrieval of the resulting 4,445 publications from the Open Access PubMed Central, the licence information recorded in the XML files was reviewed. Twenty-two documents did not contain a stated licence and were therefore excluded from the corpus, resulting in a final set of 4,423 documents with explicitly stated licence information. As these documents were not manually reviewed, they were annotated using the established automated pipelines and released as the CoDiet-Bronze (n = 4,423) and CoDiet-Silver (n = 4,423) corpora. The CoDiet-Gold corpus remained unchanged and continued to consist of the original manually curated document set.

All 4,423 re-distributable documents from the initial and additional searches were then annotated using the gold standard annotations obtained from the 450 public set as ‘dictionary’ to create an ‘electrum standard’ (electrum is an alloy that is a mixture of gold and silver) corpus, i.e. it was machine annotated, but only using human-verified entities. Disagreements between categories (e.g. dietary vs blood cholesterol) had their categories merged with an underscore (_), allowing the flexibility of keeping them in all categories, removing them, or using rule-based methods for disambiguation. Additionally, to remove noise from this set, as no human verification is performed, we removed entities that overlap with terms present in the nltk stopwords list. Finally, entities with fewer than three characters were removed from the dictionary, as their interpretation was found to be highly context-dependent given that these entities represent abbreviations that can differ in definitions between documents. This dataset has been released with all annotations as the CoDiet-Electrum (n = 4,423) corpus (4).

### Data Licensing and Redistribution

All redistributed documents across the CoDiet corpora permitted redistribution at the time of downloading these documents; their main text remains the intellectual property and views of the original authors. During corpus creation, retrieved publications were filtered according to their licensing conditions and only documents originating from the PubMed Central (PMC) Open Access (OA) collection or equivalent redistribution-permissive sources were retained for public release. Articles in the PMC Open Access subset are made available under Creative Commons or similar licences that permit reuse and redistribution according to publisher-defined terms, with the majority carrying machine-readable Creative Commons licences. Publications without redistribution rights were excluded from the released datasets. An archive of these documents in their original, unchanged formats is distributed under a CC BY-NC-ND license (51). Accordingly, the CoDiet-derived annotation datasets (4) are distributed under a CC BY-NC 4.0 licence through Zenodo, while preserving attribution and provenance metadata for each source document within the BioC `infons` fields, including original identifiers, retrieval source, and licensing information where available. These data do not contain any material that constitutes a derivation of original work. Only the documents that permit derivations to be redistributed are shared under a CC BY-NC-SA license (52).

### Named Entity Recognition Models

To evaluate the value of the CoDiet corpora for training named entity recognition systems, separate category-specific models were trained using the CoDiet-Bronze, CoDiet-Silver, and CoDiet-Electrum corpora. For each corpus, 13 individual NER models were trained, corresponding to the 13 entity categories defined in the CoDiet annotation schema, resulting in a total of 39 trained models.

#### Architecture and Training Settings

All models were fine-tuned from BioBERT v1.1, using the `dmis-lab/biobert-base-cased-v1.1` check-point (66). Separate models were trained for each entity category rather than using a single multi-category model. Each category-specific task had a three-label sequence-labelling using the BIO2 scheme (67), comprising the labels `B-category`, `I-category`, and `O`.

The models were trained using the same training script and hyperparameter configuration for all categories and corpus types. Tokenisation was performed using the corresponding BioBERT tokeniser with a maximum sequence length of 512 tokens. The corpora were prepared at the passage level, and passages were represented as sequences of up to 512 tokens. The preprocessing workflow ensured that passages exceeding this maximum length were divided into consecutive sequences before model training.

Models were trained using the `transformers` library (68), for a maximum of five epochs using a learning rate of 3 × 10^*−*5^, a per-device training batch size of 32, and a per-device evaluation batch size of 32. Gradient accumulation was performed over two steps, resulting in an effective batch size of 64 sequences. A linear learning-rate scheduler was used with a warm-up ratio of 0.1. Weight decay was set to 0.01, and gradient clipping was applied with a maximum gradient norm of 1.0. Model evaluation and checkpoint saving were performed after each epoch.

No corpus-specific hyperparameter optimisation or grid search was performed. Instead, the same architecture and training configuration were applied consistently across all 39 models to enable comparison of the effects of the different training corpora. For each model, the checkpoint achieving the highest validation F1-score was retained as the final model using the `load_best_model_at_end` procedure. Although model training was performed using token-level BIO2 representations, model performance was evaluated using strict entity-level matching. Under this evaluation criterion, a predicted entity was considered correct only when its boundaries and entity category exactly matched those of the corresponding reference annotation. Partially overlapping predictions were not considered correct.

#### Training and Evaluation Data

The 450 PMCIDs included in CoDiet-Gold-public were randomly split into 400 training documents and 50 validation documents, and this split was identical for all 39 models. The remaining 50 documents from CoDiet-Gold-private were used as an independent held-out test set and were not used during model training or validation.

For each training corpus, the corresponding annotations were converted into category-specific BIO2 representations. Annotations belonging to the target category were encoded using `B-category` and `I-category` labels, whereas all remaining tokens were assigned the `O` label. This procedure was independently applied to each of the 13 categories in the Bronze, Silver, and Electrum corpora.

The Bronze-trained models were trained using annotations generated by the Bronze automated annotation pipeline. The Silver-trained models used the corresponding annotations following overlap resolution and post-processing, whereas the Electrum-trained models were trained using annotations generated using the validated vocabulary and labels derived from CoDiet-Gold-public.

The same CoDiet-Gold-private test corpus was used to independently evaluate the trained NER models. In addition, the outputs of the original Bronze, Silver, and Electrum annotation pipelines were evaluated against the CoDiet-Gold-private annotations using the same held-out document set. This enabled comparison between the original automated annotation approaches and NER models trained using their resulting corpora.

#### Model Evaluation Results

The trained models were evaluated using two approaches. First, we assessed how well the fine-tuned BioBERT models reproduced the annotations of the corpus on which they were trained. Second, we compared the original annotation pipelines and the corresponding fine-tuned BioBERT models against the independent human annotations of CoDiet-Gold-private.

Table 6 presents the performance of the BioBERT models when evaluated using the corresponding Bronze, Silver, or Electrum annotations as the reference. For example, the Bronze models were evaluated against Bronze annotations, whereas the Electrum models were evaluated against Electrum annotations. This evaluation provides an indication of how consistently the annotation patterns generated by each corpus can be learned by the fine-tuned models.

**Table 6.**
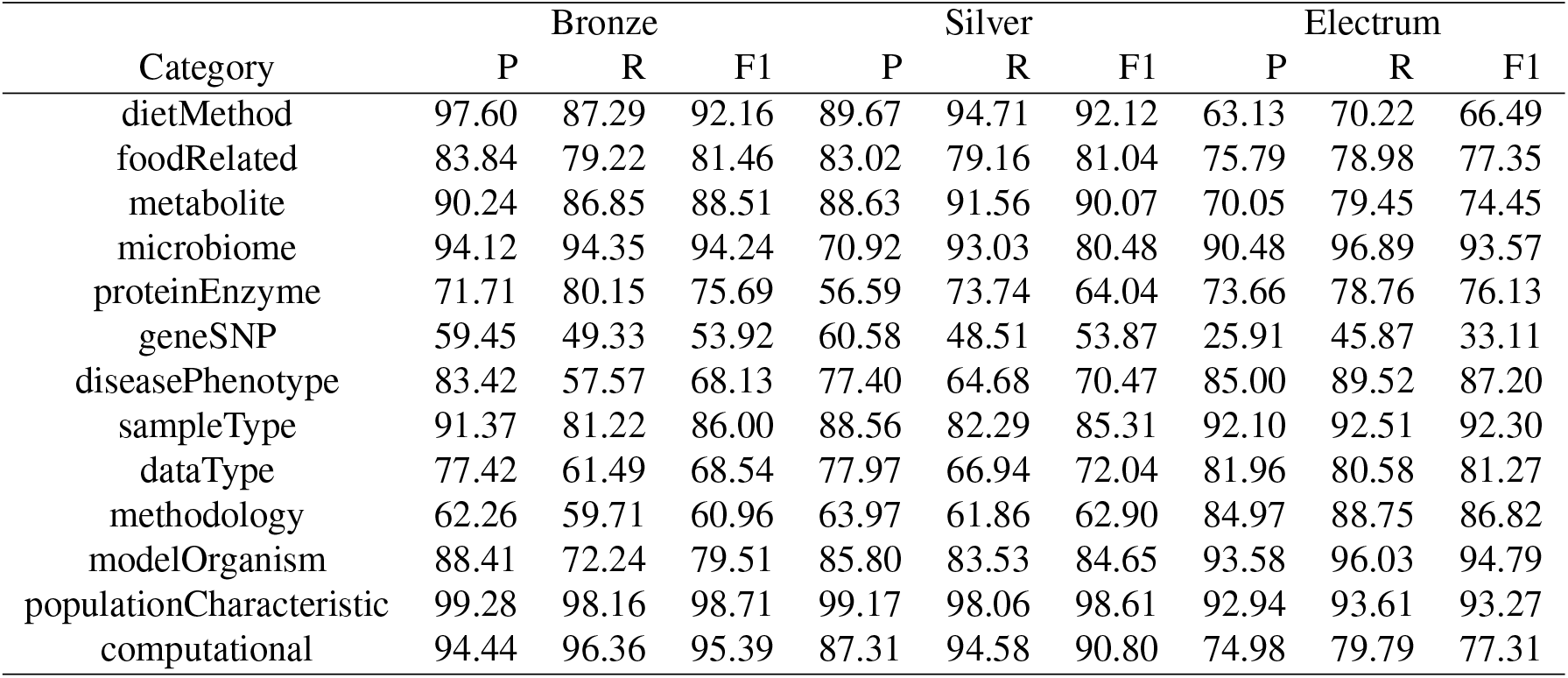
Performance of fine-tuned BioBERT models trained on 450 documents (matching to CoDiet-Gold-public) using the Bronze, Silver, and Electrum annotations and evaluated on 50 documents (matching to CoDiet-Gold-private) with Bronze, Silver, and Electrum annotations.

Performance varied between entity categories and corpus types. The highest F1-scores were generally observed for categories with more consistent annotation patterns. For example, the Bronze models achieved F1-scores above 90% for `dietMethod`, `microbiome`, `populationCharacteristic`, and `computational`. Similarly, the Electrum models achieved high F1-scores for `microbiome`, `modelOrganism`, and `populationCharacteristic`. Lower performance was observed for some categories, including `geneSNP`, `methodology`, and `proteinEnzyme`, depending on the corpus used for training.

Table 7 presents the evaluation of both the original *CoDiet-Corpus* pipeline and the corresponding fine-tuned BioBERT models against the human annotations of CoDiet-Gold-private. This evaluation provides a direct comparison between the original automated annotations and models trained using the corpora generated by the *CoDietCorpus* pipeline. The Bronze and Silver pipelines showed small variation in performance across categories. Fine-tuning BioBERT using their corresponding annotations improved performance for some categories, although this was not observed consistently across all entity types. In contrast, the Electrum pipeline generally achieved higher agreement with the human annotations across most categories.

**Table 7.** Performance of annotation pipeline and fine-tuned BioBERT models trained on 450 documents (matching to CoDiet-Gold-public) using the Bronze, Silver, and Electrum annotations and evaluated on the CoDiet-Gold-private annotations.

| Category | Model | Bronze |  |  | Silver |  |  | Electrum |  |  |
| --- | --- | --- | --- | --- | --- | --- | --- | --- | --- | --- |
|  |  | P | R | F1 | P | R | F1 | P | R | F1 |
| dietMethod | Pipeline | 2.62 | 12.67 | 4.34 | 3.04 | 12.22 | 4.86 | 87.08 | 70.14 | 77.69 |
| dietMethod | BioBERT | 2.51 | 10.86 | 4.07 | 2.56 | 10.86 | 4.14 | 69.19 | 61.99 | 65.39 |
| foodRelated | Pipeline | 37.17 | 33.33 | 35.14 | 39.99 | 31.01 | 34.93 | 86.93 | 64.35 | 73.96 |
| foodRelated | BioBERT | 40.26 | 34.12 | 36.93 | 42.60 | 31.49 | 36.21 | 81.55 | 62.92 | 71.03 |
| metabolite | Pipeline | 51.77 | 36.77 | 43.00 | 58.67 | 29.94 | 39.65 | 87.32 | 42.35 | 57.03 |
| metabolite | BioBERT | 56.11 | 38.35 | 45.56 | 59.99 | 31.63 | 41.42 | 86.37 | 47.51 | 61.30 |
| microbiome | Pipeline | 91.34 | 53.49 | 67.47 | 93.38 | 39.38 | 55.40 | 97.12 | 91.70 | 94.33 |
| microbiome | BioBERT | 92.12 | 54.08 | 68.15 | 90.70 | 50.18 | 64.62 | 96.22 | 97.28 | 96.75 |
| proteinEnzyme | Pipeline | 60.29 | 4.38 | 8.17 | 56.57 | 2.99 | 5.68 | 71.83 | 51.50 | 59.99 |
| proteinEnzyme | BioBERT | 51.32 | 4.17 | 7.71 | 41.09 | 2.83 | 5.30 | 71.99 | 55.18 | 62.47 |
| geneSNP | Pipeline | 15.91 | 46.82 | 23.75 | 18.49 | 48.45 | 26.77 | 38.07 | 13.53 | 19.98 |
| geneSNP | BioBERT | 13.89 | 33.93 | 19.72 | 15.79 | 33.12 | 21.38 | 40.67 | 25.61 | 31.43 |
| diseasePhenotype | Pipeline | 56.24 | 33.53 | 42.01 | 63.87 | 31.45 | 42.14 | 92.98 | 78.95 | 85.39 |
| diseasePhenotype | BioBERT | 74.46 | 30.64 | 43.41 | 74.46 | 30.64 | 43.41 | 84.37 | 75.45 | 79.66 |
| sampleType | Pipeline | 57.47 | 42.80 | 49.06 | 63.86 | 42.33 | 50.91 | 91.80 | 83.84 | 87.64 |
| sampleType | BioBERT | 64.70 | 42.84 | 51.55 | 69.28 | 42.67 | 52.81 | 87.57 | 80.33 | 83.8 |
| dataType | Pipeline | 20.89 | 17.79 | 19.22 | 22.86 | 18.01 | 20.14 | 91.50 | 81.30 | 86.09 |
| dataType | BioBERT | 24.48 | 16.56 | 19.76 | 24.48 | 16.56 | 19.76 | 81.47 | 71.17 | 75.97 |
| methodology | Pipeline | 10.06 | 16.29 | 12.44 | 10.82 | 16.97 | 13.22 | 90.65 | 91.46 | 91.05 |
| methodology | BioBERT | 11.79 | 18.31 | 14.34 | 11.71 | 17.75 | 14.11 | 79.85 | 84.16 | 81.95 |
| modelOrganism | Pipeline | 52.74 | 74.69 | 61.82 | 59.20 | 69.99 | 64.14 | 98.10 | 90.85 | 94.34 |
| modelOrganism | BioBERT | 66.00 | 76.38 | 70.81 | 64.70 | 74.47 | 69.23 | 95.44 | 90.71 | 93.01 |
| populationCharacteristic | Pipeline | 65.03 | 30.01 | 41.07 | 65.03 | 30.01 | 41.07 | 79.11 | 93.49 | 85.70 |
| populationCharacteristic | BioBERT | 65.77 | 30.01 | 41.22 | 65.67 | 29.97 | 41.15 | 74.86 | 89.10 | 81.36 |
| computational | Pipeline | 53.44 | 8.84 | 15.16 | 54.16 | 8.70 | 14.99 | 92.17 | 82.73 | 87.20 |
| computational | BioBERT | 51.98 | 8.77 | 15.01 | 50.77 | 8.84 | 15.05 | 77.72 | 74.23 | 75.93 |

For the Electrum corpus, the original pipeline achieved F1-scores above 85% for several categories, including `microbiome`, `diseasePhenotype`, `sampleType`, `dataType`, `methodology`, `modelOrganism`, `populationCharacteristic`, and `computational`. The fine-tuned BioBERT models produced mixed results when compared with the original

Electrum pipeline. Performance improved for some categories: `metabolite`, `microbiome`, `proteinEnzyme`, and `geneSNP`, while the original pipeline remained stronger for the rest.

The evaluation against CoDiet-Gold-private in Table 7 highlights the limitations of the automatically generated annotations. Although the pipelines achieved high agreement with the human annotations for some categories, performance varied considerably across corpus types and entity categories. The results therefore show that the annotations generated by the automated pipelines do not provide the same level of agreement with human annotations for all categories. Similarly, the performance of the fine-tuned BioBERT models varied across categories and did not consistently improve upon the original pipelines. These results provide a technical assessment of the CoDiet corpora and highlight the differences between automatically generated annotations and the independently curated human annotations.

### Usage Notes

The CoDiet datasets are intended for research in biomedical natural language processing, specifically named entity recognition involving diet, non-communicable diseases, and biomarkers. It can be used for NLP model training, evaluation, and benchmarking. Users should distinguish between the manually annotated CoDiet-Gold data and the automatically annotated datasets (CoDiet-Bronze, CoDiet-Silver, CoDiet-Electrum) as the latter may contain annotation errors but contain a larger set of documents.

## Data Availability

The CoDiet data resources are publicly available through three Zenodo releases that separately provide the original source XML files, CoDiet annotation data, and a subset of pre-combined processed full-text BioC documents.

The original unchanged XML source files obtained from Open Access PubMed Central are available as *CoDiet corpus (v*.*0*.*1) – raw XML* through Zenodo at https://doi.org/10.5281/zenodo.22097875. This release contains the source XML files corresponding to the CoDiet-Bronze, CoDiet-Silver, CoDiet-Electrum, CoDiet-Gold-public, and CoDiet-Gold-private corpora.

The corresponding CoDiet annotations are available separately as *CoDiet corpus (v*.*1*.*0) – annotations* through Zenodo at https://doi.org/10.5281/zenodo.22097879. This release contains annotation information for 4,423 documents in each of the Bronze, Silver, and Electrum corpora and for 450 documents in the Gold-public corpus.

A ready-to-use subset of processed full-text BioC JSON documents is available as *CoDiet corpus (v*.*1*.*1) – full-texts annotated* through Zenodo at https://doi.org/10.5281/zenodo.22097881. This release contains 3,965 pre-combined machine-annotated documents for each of the CoDiet-Bronze, CoDiet-Silver, and CoDiet-Electrum corpora, 409 human-annotated documents for CoDiet-Gold-public, and 46 unannotated documents for CoDiet-Gold-private.

The complete corresponding CoDiet datasets can be reconstructed from the raw XML and annotation releases using the *CoDietZenodo* workflow, available at https://github.com/omicsNLP/CoDietZenodo and archived at https://doi.org/10.5281/zenodo.22142400. This workflow converts the original XML files to BioC JSON and incorporates the corresponding CoDiet annotations into the generated documents.

The code used for corpus generation, automated annotation, and processing is available through the *CoDietCorpus* repository at https://github.com/omicsNLP/CoDietCorpus and archived at https://doi.org/10.5281/zenodo.20643594.

The CoDiet-Gold-private corpus is used for benchmarking through Codabench at https://www.codabench.org/competitions/11676/. Its evaluation annotations are not publicly released.

## Code Availability

The code and resources used to generate, process, annotate, and reproduce the CoDiet corpora are publicly available through GitHub and archived on Zenodo. Two repositories are provided to support full reproducibility of the datasets. All package and software version numbers are included in the readme documents to allow creating virtual environments to run the code.

The first repository contains the scripts and resources used for automated corpus generation and processing, including Bronze and Silver corpus creation, annotation pipelines, dictionary-based matching, and supporting processing utilities. Repository documentation provides installation instructions, software dependencies, and example commands for reproducing the released datasets.

Code repository: https://github.com/omicsNLP/CoDietCorpus

Archived release: https://doi.org/10.5281/zenodo.20643594

A second repository provides the workflow for recreating the CoDiet datasets from the original Open Access PMC XML files without modification. The workflow processes the unmodified XML files into BioC format and subsequently incorporates the released CoDiet annotations into the newly generated BioC documents. This approach enables the complete datasets to be recreated while separating the redistribution of the original source documents from the derivative annotated information.

Code repository: https://github.com/omicsNLP/CoDietZenodo

Archived release: https://doi.org/10.5281/zenodo.22142400

## Author Contributions

Conceptualization: MR, TB, and JP; Methodology: AL and JP; Software: AL, XY, TR, CCA, ADA, and JP; Validation: AL, YG, TR, DC, TB, and JP; Resources: MR, TB, and JP; Data Curation: AL, AM, SR, AC, KL, GT, MB, YG, OD, RA, NE, ADR, NB, DR, RG, DG, IT, RV, VG, MC, PP, GA, SA, NN, XY, RFC, TR, TB, and JP; Writing - Original Draft: AL and JP; Writing - Review & Editing: AL, TB, and JP; Visualization: AL and SG; Supervision: JP; Project administration: JP; Funding acquisition: GT, IT, SA, MR, TB, and JP. All authors have read and agreed to the published version of the manuscript.

## Acknowledgements

The authors thank the team of annotators for their support and help during the annotation process. A full list (ordered by annotations made, minimum of 25 articles) is given here: Alisha Mahmud, Rakesh Velmurugan, Shruti Rajendra, Katerina Loupasaki, Georgios Theodoridis, Rebeca Fernández Carrión, Maider Bizkarguenaga Uribiarte, Rubén Gil Redondo, Mercedes Caro Burgos, Ainara Cano San José, Nerea Burguera, Ricardo Diogo Alves Conde, Ángela de Diego Rodríguez, Alexander Simpson, Nieves Embade, Olga Deda, Niharika Chokkapu, Vasiliki Gkanali, Domniki Gallou, Danai Rossiou. All annotators are listed on the annotation corpus Zenodo (4). For more information see https://www.codiet.eu/. All annotators are included in the Zenodo repository.

The CoDiet study has been accomplished through work of the staff at 17 different partners. A partial listing of colleagues in the CoDiet consortium follows (alphabetical, per partner):

AUTh (Aristotelio Panepistimio Thessalonikis): Olga Begou, Olga Deda, Helen Gika, Theodore Liapikos, Danai Rossiou, Georgios Theodoridis, Dimitrios Zaikis

AZTI (Fundacion Azti - AZTI Fundazioa): Sara Arranz, Mercedes Caro, Elena Santacruz, Itziar Tueros

BRUKER (Bruker Biospin Group): Claire Cannet, Birk Schütz

CIBER (Consorcio Centro De Investigacion Biomedica En Red): Cristina Bouzas, Helmut Schroder, Antoni Sureda, Josep Tur Mari

CIC BIO (Asociacion Centro de Investigacion Cooperativa en Biociencias): Angela de Diego, Nieves Embade, Ruben Gil, Itziar Gil de la Pisa, Oscar Millet

CVUT (Ceske Vysoke Uceni Technicke V Praze): Fadwa Idlahcen, Ondrej Kuzelka, Jakub Marecek, Petr Ryšavý, He Xiaoyu

ICL (Imperial College London): Marta Archanco, Ivana Balkan, Donghee Choi, Dorota Cieslak-Jones, Aygul Dagbasi, Ben Davis, Pietro Ferraro, Gary Frost, Isabel Garcia Perez, Stamatia Giannarou, Alexandra Halbish Rayner, Monica Hill, Antoine Lain, Benny Lo, Po-Wen Lo, Heather Lodge, Baichen Lu, Siobhan Markus, George Mylonas, Jack Olney, Joram Posma, Marek Rei, Adrian Rubio, Franco Sassi, Robert Shorten, Cristina Taddei, Martina Tashkova, Jingmin Zhu

ISS (Istituto Superiore Di Sanita): Anna Ceccarelli, Valentina De Cosmi, Marco Silano

MIC (Microcaya): Ines Barretxeguren, Sabin Linaza, Natalia Zaldua

NKUA (Ethniko Kai Kapodistriako Panepistimio Athinon): Dimitrios Gunopulos, Vana Kalogeraki, Lydia Themeli, Dimitris Tomaras, Kleopatra Markou

NOT (University of Nottingham): Tim Beck, Yajie Gu, Thomas Rowlands

SCI (Sciensano): Jessie Van Kerckhove, Stephanie Vandevijvere

TAI (Tervise Arengu Instituut (National Institute for Health Development)): Diva Eeensoo, Marit Priinits

TEAGASC (TEAGASC - Agriculture and Food Development Authority): Paul Cotter, Oksana Mazur, Orla O’Sullivan

TECHNION (TECHNION - Israel Institute of Technology): Mark Davison, Mark Kozdoba, Shie Mannor, Binyamin Perets, Adaia Shiboleth

UNITN (University of Trento): Gabriel Baldanzi, Akshay Gaike, Federica Pinto, Nicola Segata

UVEG (Universitat de Valencia): Oscar Coltell Simón, Dolores Corella, Rebeca Fernandez, Carolina Ortega, Olga Portoles, Angeles Sanchis, Jose Sorli

